# Silencing of *TaCKX1* mediates expression of other *TaCKX* genes to increase grain yield in wheat

**DOI:** 10.1101/2020.01.07.897421

**Authors:** Bartosz Jabłoński, Hanna Ogonowska, Karolina Szala, Andrzej Bajguz, Wacław Orczyk, Anna Nadolska-Orczyk

**Author notes:** Corresponding author: Anna Nadolska-Orczyk. **Author contributions:** A.N-O. conceived and designed the original research plan and wrote the manuscript; B.J. performed most of the experimental work; H.O. prepared the figures and take part in data analysis; K.S. designed primers and performed comparative analysis with databases, took part in selected experiments; A.B. performed measurements of phytohormones; WO took part in planning and discussing results and helped with performing experiments.

## Abstract

*TaCKX* family genes influence development of wheat plants by specific regulation of cytokinin content in different organs. However, their detailed role is not known. The *TaCKX1*, highly and specifically expressed in developing spikes and in seedling roots, was silenced by RNAi-mediated gene silencing via *Agrobacterium* and the effect of silencing was investigated in 7 DAP spikes of T_1_ and T_2_ generations. Various levels of *TaCKX1* silencing in both generations influence different models of co-expression with other *TaCKX* genes and parameters of yield-related traits. Only a high level of silencing in T_2_ resulted in strong down-regulation of *TaCKX11 (3)*, up-regulation of *TaCKX2.1*, *2.2*, *5* and *9* (*10*), and a high yielding phenotype. This phenotype is characterised by higher spike number, grain number and grain yield, as well as slightly higher mass of seedling roots, but lower thousand grain weight (TGW) and slightly lower spike length. Content of most of cytokinin forms in 7 DAP spikes of silenced T_2_ lines increased from 40 to 76% compared to the non-silenced control. The CKs cross talk with other phytohormones.

Each of the tested yield-related traits is regulated by various up- or down-regulated *TaCKX* genes and phytohormones. Unexpectedly, increased expression of *TaCKX2.1* in silent for *TaCKX1* T_2_ plants up-regulated trans- and cis-zeatin and trans-zeatin glucosides, determining lower TGW and chlorophyll content in flag leaves but higher grain yield. The coordinated effect of *TaCKX1* silencing on expression of other *TaCKX* genes, phytohormone levels in 7 DAP spikes and yield-related traits in silenced T_2_ lines is presented.

**One-sentence summary:** Different levels of *TaCKX1* silencing influence various models of coordinated expression of *TaCKX* genes and phytohormone levels in 7 DAP spikes, as well as yield parameters.

## Introduction

Wheat (*Triticum aestivum* L.) is the third economically most important crop in the world after corn and rice and probably the most important in moderate climates. It provides approximately 20% of human calories and protein (Reynolds and Braun, 2019). The large genome of this high-yielding species, composed of three (AABBDD) genomes, has been very challenging for improving traits (Borrill et al., 2019). However, it might be a great reservoir for further increase of grain productivity (Nadolska-Orczyk et al., 2017). Continuous increase of wheat production is necessary to feed the rapidly growing world population (Foley et al., 2011) http://iwyp.org/). Biotechnological tools implemented in the process of increasing wheat productivity are expected to be beneficial.

Cytokinins (CKs) are important regulators of plant growth and development, influencing many agriculturally important processes (reviewed in Kieber and Schaller, 2018). Most of the CKs positively regulate cell division and organization of shoot stem cell centres as well as stimulating the endocycle in roots by auxin-dependent or auxin-independent mechanisms (Schaller et al., 2014). CKs are also involved in regulation of various developmental and physiological processes including size and structure during leaf development (Skalak et al., 2019), delay of senescence (Gan and Amasino, 1997; Lara et al., 2004), apical dominance (Tanaka et al., 2006), root proliferation (Werner et al., 2001; Werner et al., 2003). This regulation might occur at the posttranscriptional and/or posttranslational level (Cerny et al., 2011; Kim et al., 2012) or by modulation of context-dependent chromatin accessibility (Potter et al., 2018). CKs modulate expression of other genes involved in the control of various processes including meristem activity, hormonal cross talk, nutrient acquisition, and various stress responses (Brenner et al., 2012). There is growing evidence on their key role in seed yield regulation (reviewed by Jameson and Song, 2016). In cereals and grasses an increased content of CKs has been reported to positively affect sink potential in developing grains (Liu et al., 2013), maintain leaf chlorophyll status during plant senescence (Zhang et al., 2016) and grain filling (Panda et al., 2018). Moreover, in wheat the CKs take part in regulation of seed dormancy (Chitnis et al., 2014).

The majority of naturally occurring CKs in plants belong to isoprenoid cytokinins grouping *N^6^*-(12-isopentenyl) adenine (iP), *trans*-zeatin (tZ), *cis*-zeatin (cZ), and dihydrozeatin (DZ) derived from tRNA degradation or from isopentenylation of free adenine nucleosides catalysed by isopentenyltransferase (IPT) or tRNA-IPT. The second, smaller group comprise N6-aromatic CKs, represented by benzyladenine (BA) (Sakakibara, 2006). To better characterize their physiological role, CKs are classified into free-base active forms as tZ, cZ and iP, translocation forms (nucleosides) as tZ-ribosides (tZR), which exhibit a low level of activity, and sugar conjugates (*O*-glucosides), which are storage and inactivated forms (Sakakibara, 2006; Bajguz and Piotrowska, 2009).

CKs function as local or long-distance regulatory signals, but the mechanisms of their precise spatial and temporal control are still largely unknown (Brandizzi, 2019). They are produced in roots as well as in various sites of aerial part of plants (Kudo et al., 2010). The level of CKs in respective cells and tissues is dependent on many processes including biosynthesis, metabolism, activation, transport, and signal transduction. Active CKs can be metabolized via oxidation by cytokinin oxidase/dehydrogenase (CKX) or by activity of glycosyltransferases. Many reports have demonstrated that the irreversible degradation step by CKX enzyme plays an important role in regulation of cytokinin level in some cereals: maize (Brugiè re N, 2003), rice (Ashikari et al., 2005), barley (Zalewski et al., 2010; Zalewski et al., 2014) and wheat (Song et al., 2012; Zhang et al., 2012; Ogonowska et al., 2019).

The level of CKs might be also regulated by transport through the plant. tZ-type CKs are mainly synthesized in roots and transported apoplastically to shoots, which promote the growth of the above-ground parts of the plant (Beveridge et al., 1997; Hirose et al., 2008). In contrast, the iP and cZ type CKs are the major forms found in phloem and are translocated from shoots to roots (Corbesier et al., 2003; Hirose et al., 2008).

The *CKX* gene families in plants show different numbers of genes and various expression patterns, which are tissue- and organ-specific, suggesting gene-specific functions. Specificity of expression of 11 *TaCKX* in developing wheat plants were assigned to four groups: highly specific to leaves, specific to developing spikes and inflorescences, highly specific to roots and expressed in various levels through all the organs tested (Ogonowska et al., 2019). The *TaCKX* genes co-operated inside and among organs. Their role in plant productivity has been described in many plants including model plants and some cereals. Knock-out mutation or silencing by RNAi of *OsCKX2* in rice significantly increased grain number (Ashikari et al., 2005). The same effect of elevated grain number, spike number and yield was reported for RNAi-silenced *HvCKX1* in barley (Zalewski et al., 2010; Zalewski et al., 2012; Zalewski et al., 2014) and repeated for the same gene under field conditions (Holubova et al., 2018). Moreover, significantly increased grain number per spike was found as the effect of the *TaCKX2.4* gene silenced by RNAi (Li et al., 2018). Knock-out mutation of *HvCKX1* by CRISPR/Cas9 editing had a limited effect on yield productivity (Gasparis et al., 2019) or no yield data were supplied (Holubova et al., 2018). However, the results were contradictory for root growth. In the first report, by Gasparis et al. (2019), knock-out ckx1 mutants showed significantly decreased CKX enzyme activity in young spikes and 10-day old roots, which corresponded to greater root length, increased surface area, and greater numbers of root hairs. In the second report, by Holubova et al. (2018), root growth in *Hvckx1* mutant plants was reduced, which was similar to knock-out mutants of ckx3 showing smaller roots (Gasparis et al., 2019). The role of other *TaCKX* genes in wheat was analysed based on natural *TaCKX* variation. Haplotype variants of *TaCKX6a02* and *TaCKX6-D1* were related to higher filling rate and grain size (Zhang et al., 2012; Lu et al., 2015). QTL found in recombinant inbred lines containing a higher copy number of *TaCKX4* was associated with higher chlorophyll content and grain size (Chang et al., 2015). To arrange the numbering of *TaCKX* family genes, a new annotation for the first two was suggested by Ogonowska et al. (2019) based on the Ensembl Plants database (Kersey et al., 2018) and phylogenetic analysis. *TaCKX6a02* was annotated as *TaCKX2.1*, *TaCKX6-D1* (JQ797673) was annotated as *TaCKX2.2* and *TaCKX2.4* was annotated as *TaCKX2.2*. Annotations for these genes were maintained in the recently published review on the *TaCKX* (Chen et al. 2019), however tested in this research *TaCKX10* was renamed as *TaCKX9* and *TaCKX3* was renamed as *TaCKX11*. Newly revised by Chen et al. (2019) naming is applied and former names are in brackets.

Due to the size and complexity of the wheat genomes the knowledge about the role of *TaCKX* genes, containing homoeologues from three genomes, is more difficult to obtain, because of the limited number of natural mutants. Most homoeologues genes are expected to have overlapping functions (Uauy, 2006) therefore the effect of gene mutations might be masked by the other genomes. On average, ∼70% of wheat homoeologue triads/triplets (composed of A, B, and D genome copies) showed balanced expression patterns/the same co-expression modules (Ramirez-Gonzalez et al., 2018; Takahagi et al., 2018). One solution to silence all of them is to apply RNAi-mediated gene silencing, which allowed silencing all of the homoeologues. Moreover, this tool made it possible to obtain a number of lines with different levels of silencing, which in the case of genes coding proteins of key importance for life (when lack of protein was correlated with lethality) gave a possibility to regenerate plants for analysis (Travella et al., 2006). Introduction of a silencing cassette by stable transformation results in a stable, and inherited to T_4_, effect of silencing (Gasparis et al., 2011; Zalewski et al., 2014). The applicability of *Agrobacterium*-mediated transformation compared to a biolistic one for gene silencing of the developmentally regulated gene *HvCKX10 (2)* was proved to be reliable (Zalewski et al., 2012).

We present the first report on the role of *TaCKX1* in co-regulation of expression of other *TaCKX* genes, phytohormone content and their joint participation in regulation of yield-related traits in wheat. *TaCKX1*, silenced by a hpRNA type of vector via *Agrobacterium*, showed different levels of silencing in T_1_ and T_2_, which has been related to different models of gene action. The coordinated effect of *TaCKX1* silencing in 7 DAP spikes of T_2_ plants on expression of other *TaCKX* genes, phytohormone levels as well as phenotype based on ratio indicators is presented. Moreover, models of regulation of phytohormone levels and phenotypic traits by coordinated expression of *TaCKX* genes based on correlation coefficients (cc) in non-silenced and silenced wheat plants are proposed.

## Results

### Expression levels of silenced *TaCKX1* in segregating T_1_ and T_2_ plants

Expression levels of *TaCKX1* were measured in 44 segregating T_1_ plants from 8 T_0_ PCR+ lines. In 14 T_1_ plants relative expression (related to the control = 1.00) ranged from 0.39 to 0.88 with the mean of 0.67 (±0.14). In 30 T_1_ plants relative expression ranged from 0.90 to 1.52 with the mean of 1.16 (±0.18) (Fig. 1 A). The proportion of silenced to non-silenced plants changed in the T_2_ generation. There were 42 silenced from 0.24 to 0.88 plants with the mean of 0.54 (±0.14) and 20 non-silenced plants. Eight of them, with low relative expression ranging from 0.24 to 0.40 (mean 0.33 ±0.14), and representing different T_1_ lines, were selected for further analysis.

**Fig. 1.**
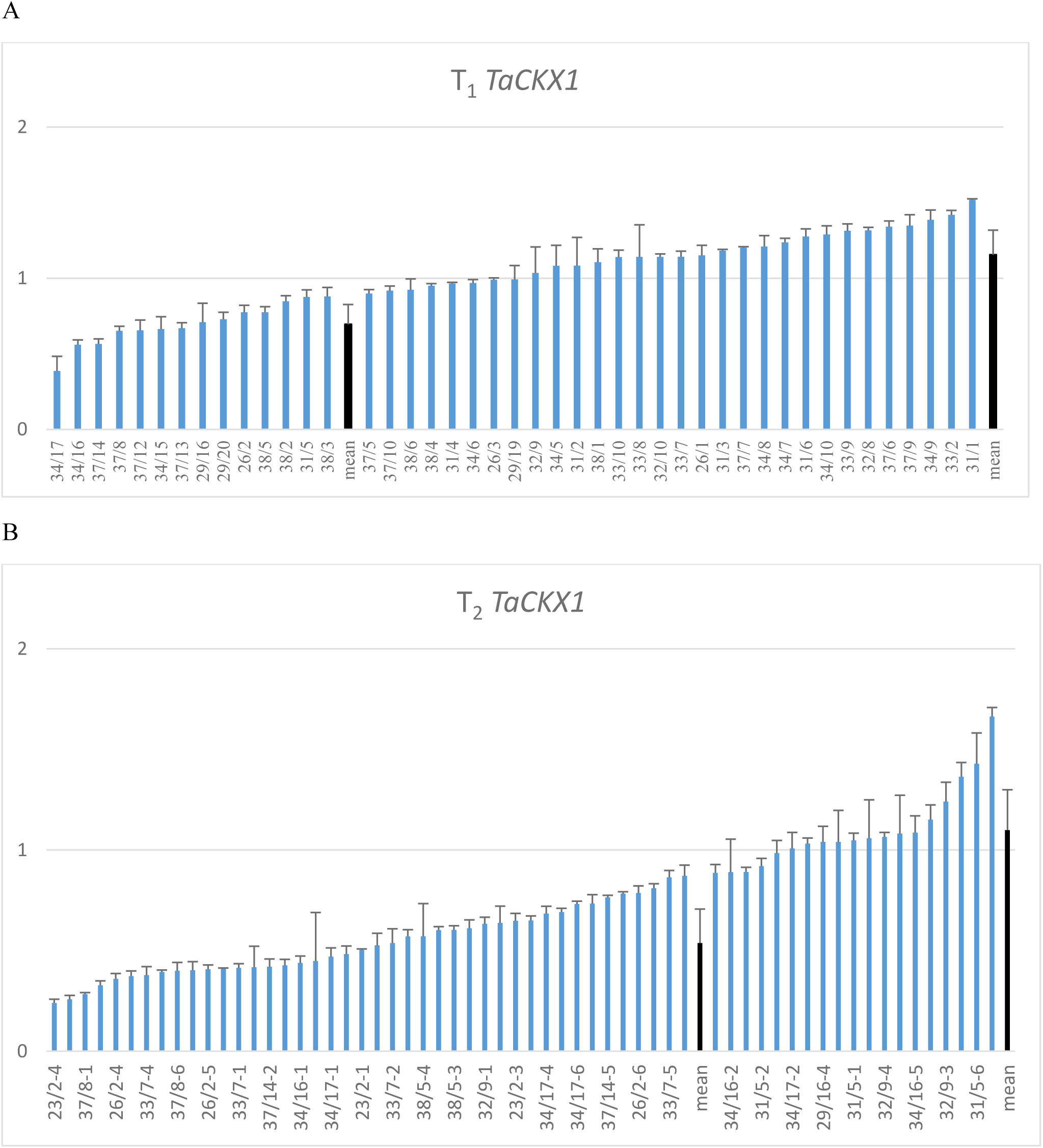
A, B. Relative expression level of silenced *TaCKX1* in segregating T_1_ (A) and T_2_ (B) plants. The level of expression is related to the control set as 1.00.

### Co-expression of silenced *TaCKX1* with other *TaCKX* genes in T_1_ and T_2_ and CKX enzyme activity

Mean relative expression of *TaCKX1* in the selected 8 lines was 0.67 in T_1_ and was decreased to 0.33 in T_2_ (Fig. 2). Similarly, in the case of *TaCKX11* (*3*) related gene expression was 0.81 in T_1_ and was decreased to 0.34 in T_2_. Relative expression levels of *TaCKX2.2* and *TaCKX9 (10)* were decreased in T_1_ to 0.51 and 0.39 and increased in T_2_ slightly above the control level, to 1.08 and to 1.10 respectively. Mean relative values for *TaCKX2.1* were similar to control in T_1_ (1.05) and slightly increased in T_2_ (1.17). Relative expression of *TaCKX5*, which was in T_1_ below the control level (0.84), was significantly increased to 1.82 in T_2_. The relative values of CKX enzyme activity in both generations were around the control, 1.00.

**Fig. 2.**
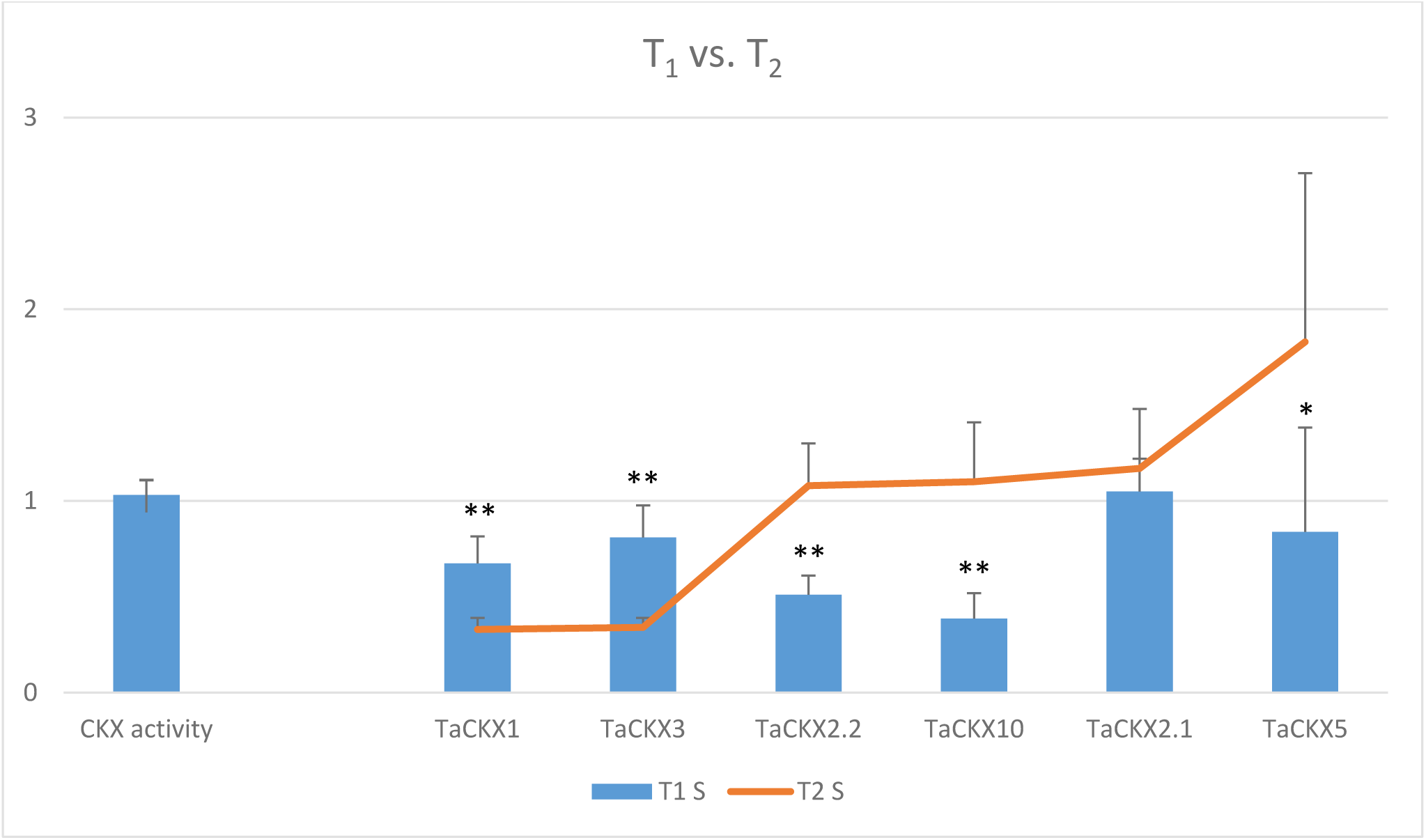
Comparison of means of relative CKX enzyme activity and selected gene expression levels in T_1_ (bars) and T_2_ (line) generation of silenced lines. * - significant at p<0.05; ** - significant at p<0.01.

The effect of *TaCKX1* silencing on the levels of expression of selected *TaCKX* genes is presented by the expression ratio indicator (Table 1), which is a quotient of the mean relative value in silent per mean relative value in non-silent, control plants. In the case of *TaCKX1* and *TaCKX11 (3)*, the ratio indicator, significantly decreased in T_1_, was strongly decreased in T_2_. The value of the ratio indicator for *TaCKX2.2* was not changed in T_1_ compared to the control and was only slightly decreased in T_2_. The expression ratio indicator of *TaCKX9 (10)*, strongly decreased to 0.59 in T_1_, rose above the control level (1.15) in T_2_. Already high in T_1_, the expression ratio indicator for *TaCKX2.1* (1.22) increased to 1.32 in T_2_.

**Table 1.** Effect of *TaCKX1* silencing on expression levels of selected *TaCKX* genes presented by expression ratio indicator (mean value in silent/mean value in non-silent, control plants) in T_1_ and T_2_ generations.

| Expression ratio indicator | T<sub>1</sub> (SD) | T<sub>2</sub> (SD) | Effect of *TaCKX1* silencing<br>T<sub>1</sub> / T<sub>2</sub> |
| --- | --- | --- | --- |
| *TaCKX1* * | 0.58 (0.12) | 0.28 (0.05) | decreased / strongly decreased |
| *TaCKX11* (3) | 0.80 | 0.36 | decreased / strongly decreased |
| *TaCKX2.2* | 1.08 | 0.98 | slightly increased / similar |
| *TaCKX9* (10) | 0.59 | 1.15 | strongly decreased / slightly increased |
| *TaCKX2.1* | 1.22 | 1.32 | increased / increased |
| *TaCKX5* | 1.00 | 1.08 | the same / similar |
| CKX activity | 1.01 | 0.99 | the same / the same |

Phenotype ratio indicator (mean value in silent/mean value in non-silent, control plants) for CKX enzyme activity was 1.01 in T_1_ and 0.99 in T_2_. Enzyme activity was not affected by the level of *TaCKX1* expression/silencing.

In T_1_ segregating plants CKX enzyme activity significantly correlated with spike length (0.51; n=16) and grain weight (0.50; n=16), but in T_2_ these correlations were not significant.

### Influence of *TaCKX1* silencing on phenotypic traits and chlorophyll content in flag leaves of T_1_ and T_2_ plants

The values of phenotypic traits in T_1_ plants with slightly decreased relative expression of *TaCKX1* (0.67 ±0.14) compared to control plants (1.00) were on the same level in the case of plant height and lower for number of spikes, spike length, grain number, and grain yield (Table S2). Higher values were obtained for TGW. Data for chlorophyll content measured by SPAD in the first spike and the next spikes were similar. All these differences were not significant. Opposite results were obtained for some traits in T_2_ plants with highly silent *TaCKX1* (0.33 ±0.06) compared to the control (1.00) (Table S3). Silent T_2_ plants were substantially smaller, had a higher number of spikes, number of grains, grain yield, seedling root weight and SPAD values for the next spikes. TGW and spike length were significantly lower than in control plants.

These differences between the slightly silenced T_1_ and highly silent T_2_ generation are expressed by comparison of ratio indicators of phenotypic traits in both generations (Fig. 3). There were no changes in plant height (cm), TGW or spike length in T_1_ plants compared to the control; however, these values were respectively 7%, 10% and 25% lower in T_2_ plants. Opposite phenotype ratio indicators for number of spikes per plant and number of grains per plant were about 21% and 30% lower in T_1_ and 57% and 29% higher in T_2_. These differences for spike number, grain number and TGW were significant.

**Fig. 3.**
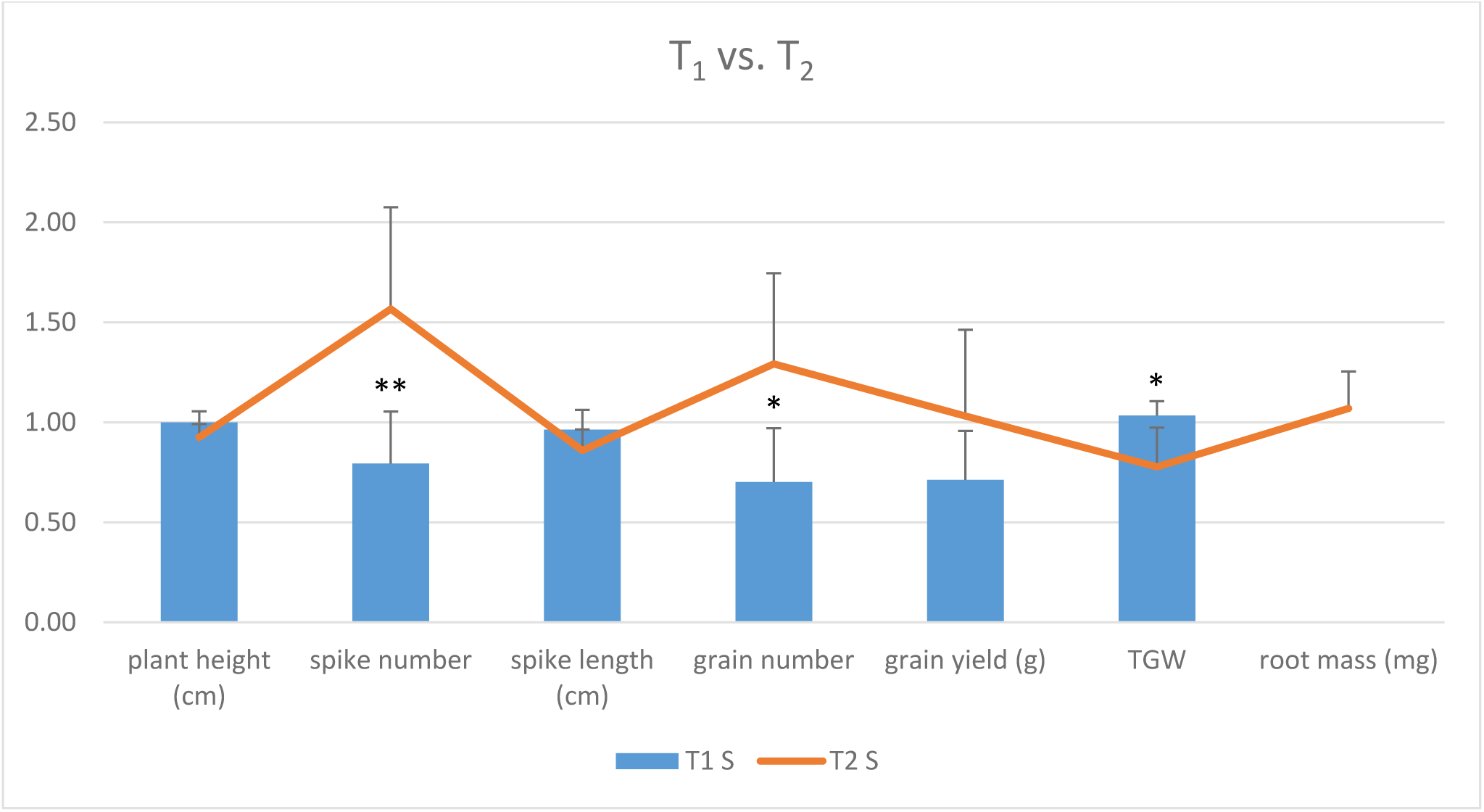
Comparison of phenotypic effect of silencing of *TaCKX1* in T_1_ and T_2_ generations based on ratio indicators. * - significant at p<0.05; ** - significant at p<0.01.

The levels of expression of *TaCKX1* in 7 DAP spikes of all T_1_ significantly correlated with number of grains, grain weight, spike length and spike number (0.47, 0.39, 0.42 and 0.33 respectively; n=42) and grain weight correlated with enzyme activity (0.33; n=42). The *TaCKX9 (10)* expression level significantly correlated with grain number (0.51; n=16).

Correlation coefficients among expression of all tested *TaCKX* genes and enzyme activity, and phenotypic traits in non-silent and highly silent T_2_ are included in Table S4 A, B. All these correlations are graphically presented in Fig. 6 A-H and described in section 6 of results.

### Phytohormone content in 7 DAP spikes of T_2_

tZOGs, which were mainly composed of tZ-*O*-glucoside (tZOG) and tZ-9-glucoside-*O*-glucoside (tZ9GOG), were the most abundant cytokinin group in 7 DAP spikes (Fig. 4 A). Their mean content in control plants was 861.18 and 452.43 ng/g biomass and in silent T_2_ was 953.86 and 974.1 ng/g biomass respectively. The second most abundant was tZ with the level of 119 ng/g biomass in the control and about twice as high in silent T_2_ (230). The next was *cis*-zeatin *O*-glucoside cZOG, which was more abundant in the control than the groups of silent plants, and the content was 63.48 and 28.68 ng/g biomass respectively. Opposite to cZOG but similar to tZ, cZ was less abundant in control (29.03 ng/g biomass) but more than three times more abundant in silent (51.03 ng/g biomass) T_2_ plants. Concentration of DZGs (sum of DZ7G, DZOG and DZOGR) was higher in silent (29 ng/g biomass) than in control plants (21 ng/g biomass). Very low concentrations (below 0.5 ng/g biomass) were measured for iP and BA. Concentration of IAA was comparable in control and in silent plants (14.26 and 14.88 ng/g biomass respectively). In the case of ABA, the low concentration in the control was slightly decreased in silent plants (2.61 and 2.29 ng/g biomass respectively).

**Fig. 4.**
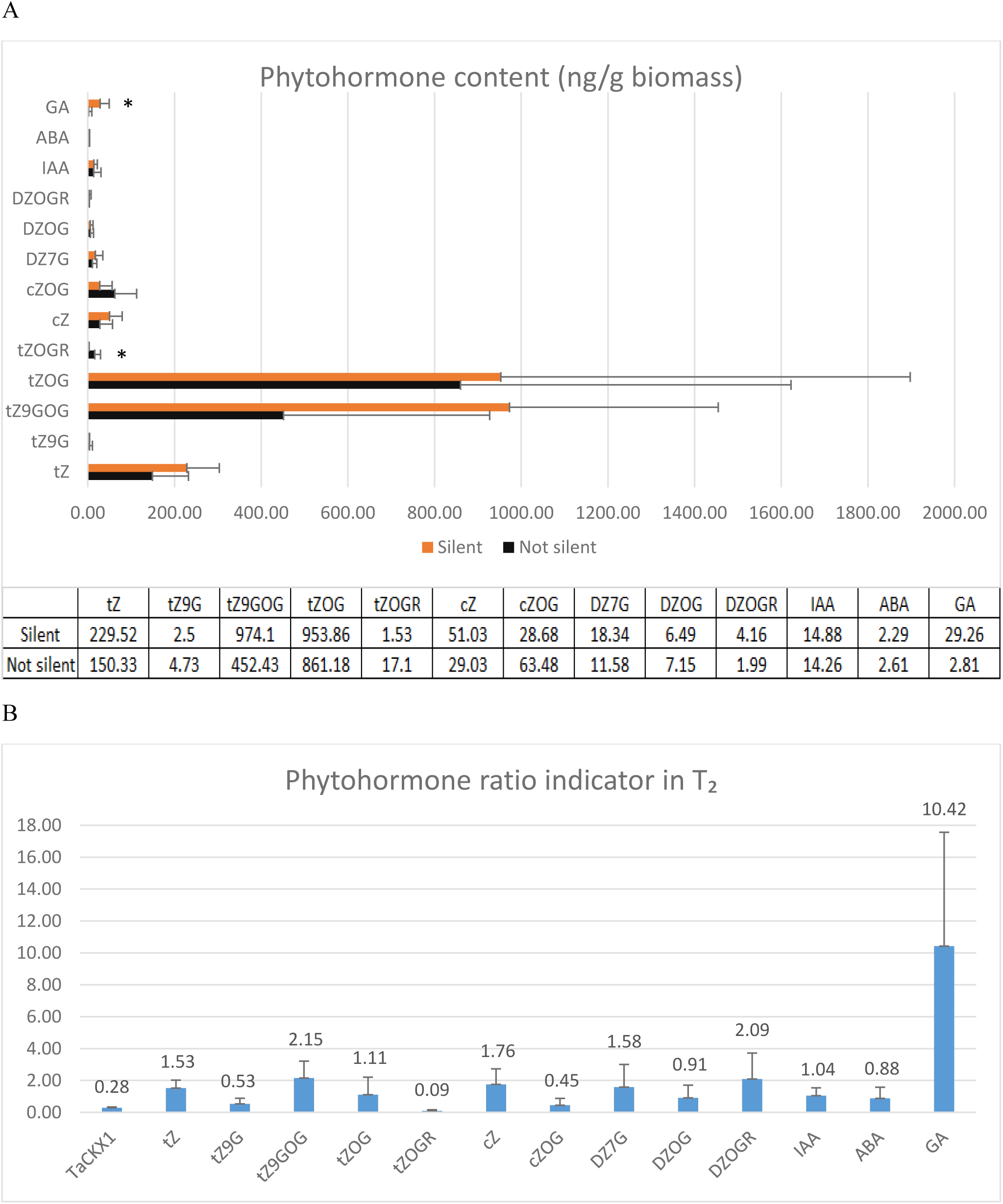
A, B. Phytohormone content (ng/g biomass) measured in the group of control and silent T_2_ plants (A). Phytohormone ratio indicators (mean value in silent per mean value in non-silent, control plants) in silent T_2_ plants (B). * - significant at p<0.05. Trace amounts (≤ 1.00 ng/g biomass): tZR, tZ7G, cZ9G, cZOGR, DZ9G, iP, iP7G, BA. Not detected: cZR, DZ, DZR, iPR, IBA, IPA, NAA, PAA.

Concentration of GA was increased from 2.81 ng/g biomass in the control to 29.26 ng/g biomass in silent plants, which was more than a 10-fold increase.

Most of the phytohormone ratio indicators in the group of six silent T_2_ plants (Fig. 4 B) were much higher than in control plants. There were the following cytokinins: tZ (1.53), tZ9GOG (2.15), tZOG (1.11), cZ (1.76), sum of DZGs (1.40) and iP (1.32). The ratio indicators for some of them were significantly lower, as in the case of BA (0.27) and cZOG (0.45). Similar values were observed for IAA, slightly lower for ABA (0.88), but much higher for GA (10.42).

### Coordinated effect of *TaCKX1* silencing on expression of other *TaCKX* genes and phytohormone level in 7 DAP spikes as well as phenotype in T_2_

A graphic presentation of the coordinated effect of *TaCKX1* silencing on expression of other *TaCKX* genes and phytohormone levels in 7 DAP spikes as well as the phenotype of T_2_ plants is presented in Fig. 5. The significant decrease of expression of *TaCKX1* was coordinated with the significant decrease of *TaCKX11 (3)*, which presumably resulted in a significant increase of most CKs: tZ, tZGs, cZ, DZGs, iP as well as GA. The increased phytohormone level in the first 7 DAP spike positively influenced traits such as spike number and grain number, reaching the ratio indicators 1.57 and 1.29, respectively, and negatively TGW (0.78), spike length (0.86), plant height (0.93) and flag leaf senescence (0.95). Opposite data were obtained for *TaCKX2.1* and *TaCKX9 (10)*, which showed increased expression in silenced, 7 DAP spikes (1.32 and 1.15 respectively). This might have influenced the decreased ratio indicators for phytohormones – cZOG (0.45), BA (0.27) and ABA content (0.88) – and slightly increased ratio indicators for yield-related traits: root weight and grain yield (1.07 and 1.03 respectively). Expression ratio indicators for *TaCKX5* and *TaCKX2.2* were both close to 1.00, but their expression significantly increased compared to T_1_ and positively correlated with the expression of *TaCKX2.1* and *TaCKX9 (10)* respectively.

**Fig. 5.**
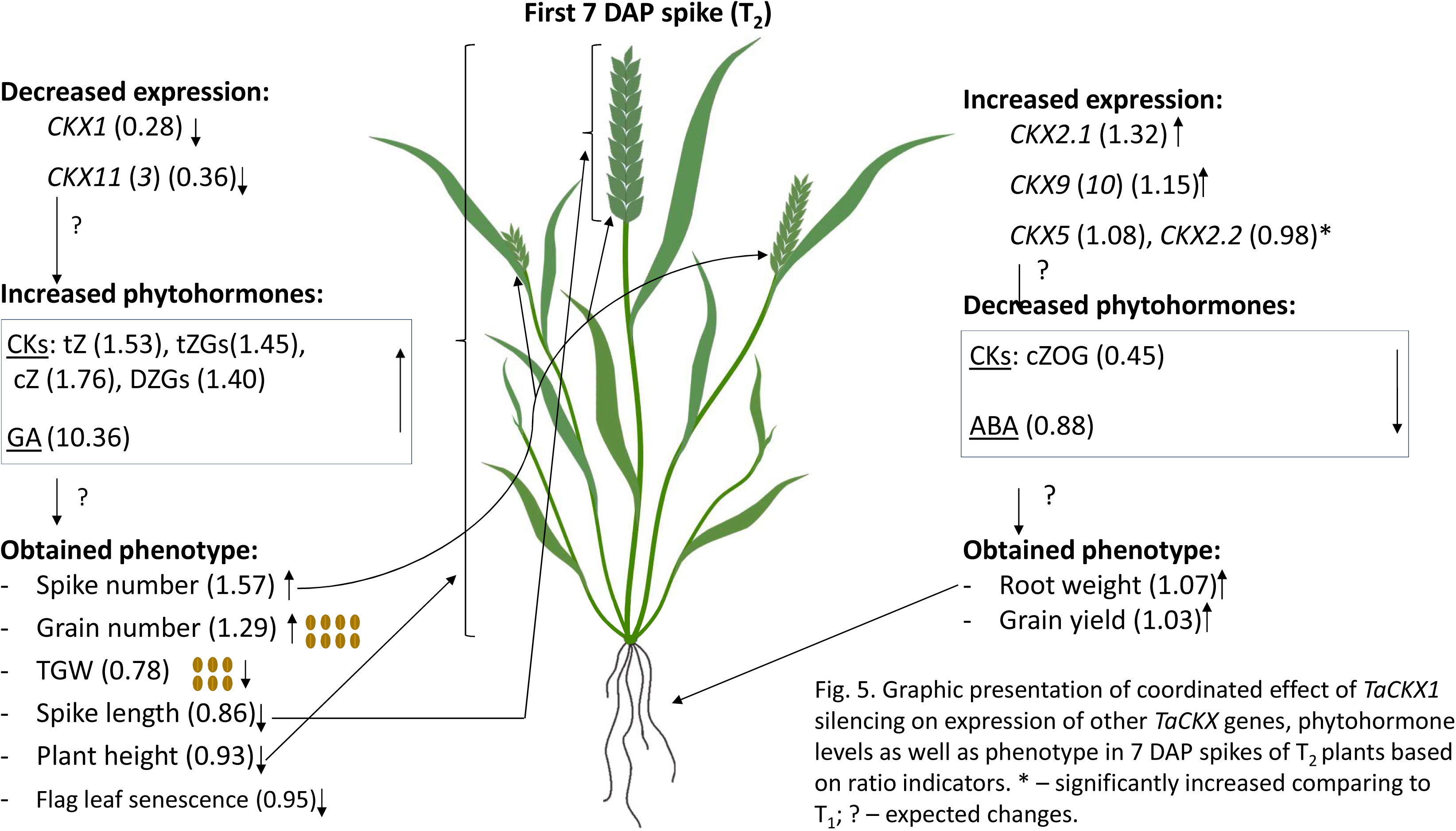
Graphic presentation of coordinated effect of *TaCKX1* silencing on expression of other *TaCKX* genes, phytohormone levels as well as phenotype in 7 DAP spikes of T_2_ plants based on ratio indicators. * – significantly increased comparing to T_1_; ? – expected changes.

### Models of co-regulation of phytohormone levels and phenotype traits by coordinated expression of *TaCKX* genes in non-silenced and silenced T_2_ plants

Two different models of co-regulation of *TaCKX* expression and phenotypic traits in non-silenced and silenced plants of the T_2_ generation are proposed (Fig. 6 A-H) based on correlation coefficients (Table S4 A, B).

Plant height (Fig. 6 A). There was no correlation between plant height and expression values of any *TaCKX* expressed in 7 DAP spikes of non-silent as well as silent plants. In the first group of plants this trait negatively correlated with BA and positively with IAA and GA content. By contrast, in silent plants the values of plant height were negatively correlated with growing concentration of tZ and tZGs, which resulted in a smaller plant phenotype.

**Fig. 6.**
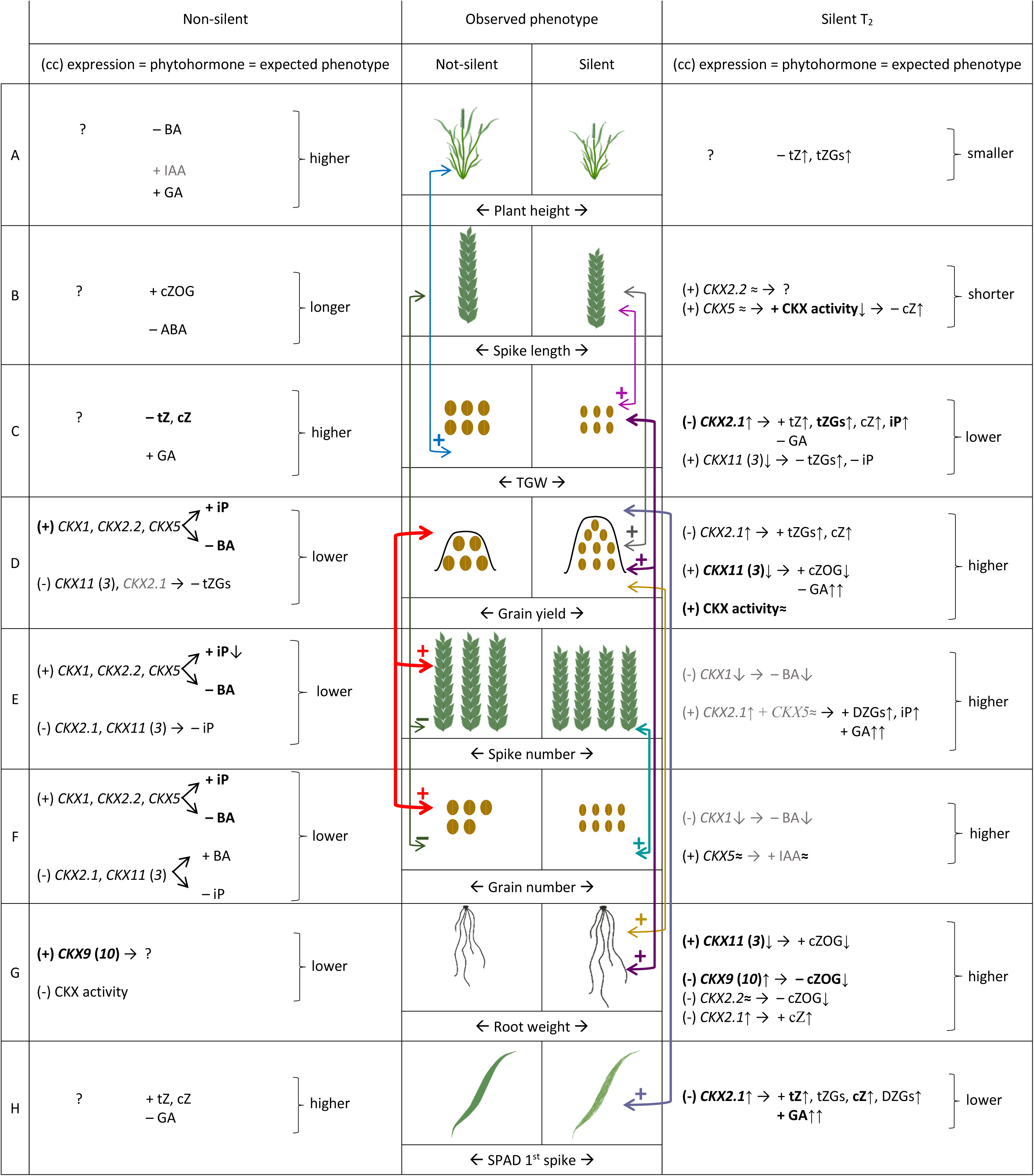
A-H. Models of regulation of phytohormone levels and phenotypic traits by coordinated expression of *TaCKX* genes based on correlation coefficients (cc) in non-silenced and silenced wheat plants. (cc) – correlation coefficient between expression and trait; bold – strong, significant correlations at p ≤ 0.05 (above 0.82); gray – from 0.5 to 0.6 ↑ – increased; ↓ – decreased; cZOG – cZ-0-glucoside (inactive); tZGs. – mainly tZ-0-glucoside + tZ-9-glucoside-0-glucoside.

Spike length (Fig. 6 B) in non-silent plants was positively correlated with cZOG, and negatively with ABA content. These correlations determined longer spikes and the trait negatively correlated with spike number and grain number. A strong positive correlation between CKX activity and spike length was noted in silent plants. The values of enzyme activity correlated positively with slightly increased *TaCKX5* expression, which negatively correlated with increasing content of cZ. Spike length in silent plants was positively correlated with grain yield.

TGW (Fig. 6 C). There was no correlation of TGW with expression of any *TaCKX* expressed in 7 DAP spikes of non-silent plants; however, the trait was strongly negatively correlated with tZ and cZ content and positively with GA. The grains in this group of plants were larger and TGW higher. By contrast, in silent plants there was a strong negative correlation of the trait with growing expression of *TaCKX2.1*, which positively regulated tZ, tZGs, cZ and iP, but negatively GA content. Moreover, the values of expression of down-regulated *TaCKX11 (3)* negatively correlated with growing content of tZGs and iP and positively with the trait. Altogether it resulted in lower TGW compared to non-silenced plants. The trait in silent plants was strongly and positively correlated with grain yield (0.82) and root weight (0.77). Grain yield (Fig. 6 D). Expression levels of *TaCKX1*, *TaCKX2.2* and *TaCKX5* in non-silent plants positively correlated with iP and negatively with BA content. However, expression of *TaCKX11 (3)* and *TaCKX2.1* negatively regulates tZGs. Altogether it resulted in lower grain yield comparing to silenced plants, and the trait was strongly positively correlated with spike number (0.93) and grain number (0.99). Increasing expression of *TaCKX2.1* positively correlated with growing content of tZGs and cZ and negatively with the trait in silent plants. Decreasing expression of *TaCKX11 (3)*, which was positively correlated with decreased cZOG content and negatively with GA content, positively correlated with the trait. Moreover, a positive correlation was observed between CKX activity and grain yield in this group of plants, which was higher than in non-silent plants. The trait was strongly correlated with TGW (0.82) and root weight (0.66).

Spike number (Fig. 6 E) and grain number (Fig. 6 F) in non-silenced plants were positively regulated by *TaCKX1*, *TaCKX2.2* and *TaCKX5*, and their expression was positively correlated with iP and negatively with BA. On the other hand, expression levels of *TaCKX2.1* plus *TaCKX11 (3)* were negatively correlated with the traits as well as with iP and positively with BA. Both groups of genes finally affected lower spike and grain number in non-silent comparing to silent plants and were strongly and positively correlated with each other (0.91) and grain yield (0.93 and 0.99 respectively). In silent plants decreasing expression of *TaCKX1* is negatively correlated with both spike and grain number and the gene negatively regulates decreasing BA content. In the case of grain number, the main player positively correlated with the trait is *TaCKX5*, increased expression of which was correlated with slightly higher IAA content, which resulted in higher grain number. Spike number is also positively regulated by *TaCKX5* co-expressed with *TaCKX2.1*, and both genes were positively correlated with growing CKs, DZGs and iP as well as GA, determining higher spike number. Both traits are highly correlated (0.88) with each other.

Seedling root weight (Fig. 6 G). There was strong, positive correlation between *TaCKX9 (10)* expression in 7 DAP spikes and seedling root weight in non-silenced plants, although the level of expression of the gene did not correlate with any phytohormone content. Moreover, CKX activity negatively correlated with the trait, which finally resulted in lower root weight. Decreasing expression of *TaCKX11 (3)* in the case of silent plants was positively correlated with decreasing content of cZOG and strongly positively correlated with the trait. Increasing expression levels of *TaCKX9 (10)* plus *TaCKX2.2* negatively correlated with decreasing content of cZOG and root weight.

Chlorophyll content measured by SPAD in flag leaves of first spikes (Fig. 6 H). There was no correlation between expression level of any *TaCKX* measured in 7 DAP spikes of non-silent plants and the trait. The only correlations were between phytohormone content and the trait, positive for tZ and cZ and negative for GA, which resulted in higher SPAD values (chlorophyll content). Increasing expression of *TaCKX2.1* was strongly positively correlated with growing values of tZ, tZGs, cZ and DZGs as well as GA in silent plants. A strong negative correlation was observed between the gene expression and chlorophyll content, which means that increasing expression of *TaCKX2.1* in 7 DAP spikes results in lower chlorophyll content in silent plants.

## Discussion

First, 7 DAP spike was chosen as a research objective in wheat since decreased *HvCKX1* expression at this stage in barley resulted in higher yield due to the higher spike and grain number (Zalewski et al., 2010; Zalewski et al., 2014). The samples were taken from the middle part of the spikes, when anthesis starts, in order to ensure a similar developmental stage of spikelets for research. The 7 DAP spikes of wheat represent the middle of cell division/cell expansion stage (Gao et al., 1992; Hess et al., 2002). The 7-day old embryo is in the early stage of development, beginning to change from globular to torpedo shape, with the root pole starting to differentiate and endosperm starting to degenerate (http://bio-gromit.bio.bris.ac.uk/cerealgenomics/cgi-bin/grain3.pl).

The *TaCKX1* gene is an orthologue of *HvCKX1* and both genes are specifically expressed in developing spikes (Ogonowska et al., 2019). In our earlier research with barley we hypothesized and continue here to address the hypothesis that the level and pattern of expression of a defined *CKX* family gene might determine the specific phenotype and indicate its function (Zalewski et al., 2014).

### Various levels of *TaCKX1* silencing influence different models of co-expression with other *TaCKX* genes and parameters of yield-related traits

One third of segregating T_1_ plants showed a decreased expression level of *TaCKX1* between 12% and 61%. Further selection of T_2_ led to obtaining plants with much higher, exceeding 60%, silencing of the gene. Different levels of silencing of *TaCKX1* in T_1_ and T_2_ generate various results of co-expression with other *TaCKX* genes and plant phenotype. Slightly decreased expression of *TaCKX1* in T_1_ was correlated with slightly decreased expression of *TaCKX11 (3)* and *TaCKX5* and significantly decreased *TaCKX2.2* and *TaCKX9 (10)*. Highly decreased expression of *TaCKX1* in T_2_ was correlated with highly decreased expression of *TaCKX11 (3)* and highly increased *TaCKX5*, *TaCKX2.2* and *TaCKX9 (10)*, and expression of *TaCKX2.1* was on a similar level to the control in T_1_ and slightly increased in T_2_. Expression of *TaCKX9 (10)* was highly and significantly correlated with *TaCKX1* only in T_1_. However, a new and strong positive correlation between *TaCKX9 (10)* and *TaCKX2.2* in highly silenced T_2_ was observed. Slightly decreased co-expression of silenced *TaCKX1* together with *TaCKX11 (3)* in T_1_ and stronger in T_2_ indicate their positive co-regulation. It should be underlined that there is no homology between the sequence of *TaCKX1* used for silencing and sequences of other *TaCKX* genes tested; therefore the process of RNAi silencing was specifically addressed to *TaCKX1* silencing. It indicates that the level of silencing of the modified gene affected variable levels of expression of the other *TaCKX* genes in a co-operative process maintaining homeostasis of CKX enzyme in the research object.

Transcriptome analysis of knock-out *HvCKX1* in barley (Gasparis et al., 2019) was associated with down- or up-regulation of other *HvCKX* family genes as well. Besides, models of co-regulation of other *CKX* by highly silenced *TaCKX1* and knock-out *HvCKX1* differ between these species.

The differences in the levels of expression of *TaCKX1* and various co-expression of other *TaCKX* genes in T_1_ and T_2_ resulted in opposite phenotypic effects. Since spike number, grain number and grain yield were reduced in T_1_, representing slightly decreased *TaCKX1* expression, the same yield-related traits were significantly higher in highly silenced T_2_ plants. High-yielding phenotype occurred when highly silenced *TaCKX1* co-operated with down-regulated *TaCKX11 (3)* but up-regulated *TaCKX5*, *TaCKX2.2*, *TaCKX2.1* and *TaCKX9 (10)*. These differences showed that both levels of silencing might be helpful to better understand the function of developmentally regulated genes. Unexpectedly, changes in expression levels of co-working *TaCKX* did not result in different enzyme activity, even in highly silenced T_2_ plants. It might be explained that down-regulation of *TaCKX1* and *TaCKX11 (3)* is compensated by up-regulation of *TaCKX2.2*, *TaCKX5* and *TaCKX9 (10)*, and therefore the contribution of isozymes encoded by the genes in the general pool of CKX enzyme activity is the same. Since CKX enzymes indicate different specificities for the particular cytokinin hormone(Gajdosova et al., 2011), the cytokinin contribution and phenotypic traits of modified plants were changed accordingly, with consequent differences in the active pool of CKs influencing phenotype.

Schaller et al. (Schaller et al., 2014) listed three possible explanations of differing, positive or negative regulatory roles of CKs in the process of cell division. The first two are the presence of additional specific regulators or other signals, which have also been reported by others (Reid et al., 2016; Mao et al., 2019), and the third possibility is the level of cytokinin activity. Involvement of additional specific regulators in differentiating the regulatory roles of CKs is also plausible in our case. However, the most convincing explanation is changed levels of CK activity (concentrations), which was proved to be significantly increased in silent plants and is further discussed below.

### Co-operating effect of *TaCKX* on the level of active CKs in silenced plants

Since *TaCKX* family genes encode CKX isozymes, which specifically degrade CKs, we might expect that the consequence of decreased expression of *TaCKX* genes is a decrease of isozyme activity in the general pool of CKX enzymes, and increased content of active cytokinins in the respective organ. Therefore according to our diagram (Fig. 5), highly decreased expression of *TaCKX1* and *TaCKX11 (3)* in 7 DAP spikes is expected to result in the observed increase of most major, active forms of CKs. Already the highest content of tZGs and tZ in non-silenced plants was increased by 45-53% in silenced plants. The third was cZ, with about ¼ of the content of tZ in non-silenced and increased by 76% in silenced plants. Conversely, the higher level of cZOG in non-silenced plants was decreased to 45% in silenced plants. We documented that both tZ and cZ, which are isomers of zeatin, together with their derivatives are a major group of isoprenoid CKs in 7 DAP spikes. It has already been shown that trans-zeatin is the predominant form after anthesis (Morris et al. 1993, Hess et al. 2002), but comprehensive analysis of cytokinins during spike, spikelet, ovule and grain development has not yet been reported for wheat using LC-MS/MS (Chen et al. 2019).

The content of DZGs increased by 40% in silent compared to non-silent wheat plants, suggesting that this less known isoprenoid form of CKs might also play an important role in plant productivity. Interestingly, isoprenoid iP was represented in 7-DAP spikes of non-silent plants at very low quantities, but its content in 7 DAP spikes of silent plants was increased by 32%. A similar relationship between reduced expression of selected *CKX* family genes and cytokinin accumulation in reproductive organs has been observed in other species including *A*. *thaliana* (Bartrina et al., 2011), rice (Ashikari et al., 2005) and barley (Holubova et al., 2018), but detailed data are not comparable to our research in wheat.

The physiological significance of these isoprenoid forms is not very well known. In maize plants iP and tZ are the most abundant and are susceptible to CKX enzyme (Bilyeu et al., 2001). DZ, which generally occurs in small quantities, is biologically stable and resistant to CKX. cZ was found to be less active and more stable than tZ and iP because of its low affinity to CKX (Bilyeu et al., 2001). Also many other researchers indicate tZ as a bioactive cytokinin, whereas cZ was reported to have a weak biological impact and unknown biological role (Schafer et al., 2015). According to Gajdosova et al. (2011), cis-Z-type CKs are frequent in the developmental stages, where they are associated with limited growth, although the metabolic fate of both cZ and tZ varied between different species. In addition, cZ levels change significantly during development in maize grain, as well as in shoot and root tissues (Saleem et al., 2010; Zalabak et al., 2014). The high levels of cZ at the first developmental stage of barley spike observed by Powell et al. (Powell et al., 2013) might indicate a possible role of this form in early barley embryo development. A significant increase of active cZ was also observed in 7 DAP spikes representing early stages of embryo development in our research and was negatively correlated with some yield-related traits such as higher grain yield and root mass and shorter spike length and lower TGW (discussed further below).

The BA is represented in 7-DAP spikes of wheat at trace amounts but their content was significantly decreased in silent plants. However, their correlations with the *TaCKX* genes as well as yield-related traits of non-silenced plants indicate their importance (discussed in more detail below). Interestingly, BA was found to participate in posttranscriptional and/or posttranslational regulation of protein abundance in *Arabidopsis*, showing high specificity to shoots and roots, and affected differential regulation of hormonal homeostasis (Zd’arska et al., 2013).

### Cross talk of CKs with other phytohormones

Negative correlations between ABA content and *TaCKX2.2* and *TaCKX9 (10)* expression, and positive with *TaCKX11 (3)*, were associated with a slight decrease of ABA content in 7 DAP spikes of silenced plants. Moreover, ABA was strongly positively correlated with BA. The main auxin, IAA, remained at the same level. A ten-fold increase of GA content in silenced comparing to non-silenced plants was observed.

Such cross regulation of CKs and other plant hormones is documented in other species. In maize kernels the *CKX1* gene is up-regulated by cytokinin and ABA, and abiotic stress (Brugiére N, 2003). In tobacco altered cytokinin metabolism affected cytokinin, auxin, and ABA contents in leaves and chloroplasts (Polanska et al., 2007), which host the highest proportion of CK-regulated proteins (Cerny et al., 2013). Moreover, auxin, ABA and cytokinin are involved in the hormonal control of nitrogen acquisition and signalling (Kiba et al., 2011), which often limits plant growth and development. All four phytohormones CKs, GA, IAA and ABA were found to be involved in regulation of grain development in drought conditions (Abid et al., 2017). Moreover, in shoots, BA up-regulated the abundance of proteins involved in ABA biosynthesis and the ABA response, whereas in the roots, BA strongly up-regulated the majority of proteins in the ethylene biosynthetic pathway (Zd’arska et al., 2013). We proved that IAA, GA and ABA contents are also co-regulated by CKs in non-silenced and silenced 7 DAP spikes. Up-regulation of major CKs and down-regulation of some minor ones in silent plants influence GA, ABA and IAA content in a similar manner as in abiotic stress conditions.

### Coordinated effect of *TaCKX* gene expression on the content of CKs, other phytohormones and yield-related traits

Plant height in non-silenced plants is down-regulated by BA and up-regulated by IAA and GA content in the first 7 DAP spikes, resulting in taller plants. Oppositely, increased content of tZ and tZGs negatively correlated with the trait in silent plants, and therefore increase of these main CKs in developing spikes negatively stimulated plant height. Since expression of genes was measured in 7 DAP spikes, none of the individual genes was shown to regulate the trait.

As it was proved by Brenner and Schmulling (Brenner and Schmulling, 2012) and similarly to our results, plant height and root weight are regulated by CKs and IAA in opposite ways. It may be dependent on basipetal auxin flow in the stem, which suppresses axillary bud outgrowth, and similarly as in pea, auxin derived from a shoot apex suppresses the local level of CKs in the nodal stem through the regulation of *CKX* or *IPT* genes (Shimizu-Sato et al., 2009).

The main role in spike length seemed to be played by cZ and its derivatives. Increased content of cZOG in non-silenced plants negatively correlated with ABA, resulting in longer spikes. In silent plants the trait is positively regulated by *TaCKX2.2* together with *TaCKX5*, and the latter is a positive regulator of enzyme activity and negative of cZ content. Consequently higher content of cZ in 7 DAP spikes led to shorter spikes. cZOG found as positive regulators of longer spikes are sugar conjugates of cZ-0-glucoside, which are inactivated forms of cZ, showing metabolic stability against CKX activity (Sakakibara, 2010). The tZ-0-glucosides hyper-accumulates in *IPT* overexpressing plants (Zubko et al., 2002). 0-glucosylation of cZ is catalysed by a specific 0-glucotransferase, cisZOG1, discovered in maize (Martin et al., 2001), and this form mainly functions in the early stages of seed development.

Knowledge of function of cZ degradation pathways via the CKX enzyme is limited. Interestingly, two *Arabidopsis* genes, *CKX1* and *CKX7*, expressed in stages of active growth, were shown to have high preference for cZ (Gajdosova et al., 2011). Accordingly, overexpression of *CKX7* highly decreased levels of free cZ in *Arabidopsis* (Kollmer et al., 2014). In our case relatively low expression of *TaCKX2.2* together with *TaCKX5* led to lower CKX activity and higher cZ content.

None of the tested individual *TaCKX* genes was involved in high TGW in non-silenced plants, but a negative correlation with tZ and cZ and positive with GA was found. A significant negative correlation of *TaCKX2.1* and a positive correlation of *TaCKX11 (3)* in determining low TGW were observed in silenced plants. Unexpectedly increased expression of the first one positively influenced tZ, tZGs, cZ and iP content and negatively GA content, and the opposite was true for the second gene, resulting in lower TGW. Therefore both *TaCKX2.1* and *TaCKX11 (3)*, acting in an opposite manner, maintain homeostasis of CKX enzyme activity and co-regulate TGW in silenced plants.

Greater concentration of CKs, especially tZ, was observed during the grain filling stage of high-yielding cultivars (Powell et al., 2013). We might suppose that the observed higher concentrations of tZ and other CKs at the 7 DAP stage, which originally was a consequence of *TaCKX1* silencing, might accelerate germination of the grains, which resulted in smaller grains/lower TGW than in non-silenced plants. The silenced *TaCKX1* co-work with down-regulated *TaCKX11 (3)* in increasing CK content as well as up-regulating *TaCKX2.1*, with seems to play a regulatory role.

The involvement of GA in TGW and other traits demonstrated by us might be the effect of co-regulation of *CKX* and other gibberellin-responsive genes. One of them is the dwarfing gene *Rht12*, which can significantly reduce plant height without changing seedling vigour and substantially increase ear fertility in bread wheat (Chen et al., 2018). Similar results were obtained with allelic variants of *Ppd-D1* (chromosome 2D) and *Rht-D1* (chromosome 4D) loci, which affect some plant growth traits, e.g. leaf area and spike length (Guo et al., 2018). Fahy et al. (Fahy et al., 2018) in their research on starch accumulation suggested that final grain weight might be largely determined by developmental processes prior to grain filling.

This is in agreement with our observations, in which yield-related traits are differently regulated in two groups of plants, non-silent and silent. Therefore we might suppose that coordinated co-regulation of expression of *TaCKX* genes and related CKs takes place during whole plant and spike development and small seeds in silenced plants are determined at earlier stages.

Grain yield, which is very strongly correlated with grain and spike number in non-silent plants but with TGW in silent plants, is a more complex feature. Two groups of genes up-regulating or down-regulating grain yield in non-silent plants have been found. The first one includes *TaCKX1*, *2.2* and *5* positively regulating iP and tZGs content but negatively BA. The second comprises *TaCKX11 (3)* (and *2.1*) acting in the opposite way. We might expect that these genes, which negatively correlate with the trait, are main determinants of lower grain yield, affected by down-regulation of tZGs and BA. It is worth to mention that *TaCKX5* is highly expressed in inflorescences and leaves. Higher grain yield was positively regulated by enzyme activity and both *TaCKX11 (3)* as well as *TaCKX2.1* in silenced plants. Indeed, correlation of *TaCKX2.1* expression with the trait was negative. However, the gene positively regulated tZGs and cZ content just like for TGW, which is rather untypical for a gene encoding a CKX enzyme degrading CKs. Therefore the positive regulation of the main CK content by *TaCKX2.1* observed by us supports its role in regulation of expression of other genes rather than encoding the CKX isozyme.

As observed in barley cultivars, changes in cytokinin form and concentration in developing kernels correspond with variation in yield (Powell et al., 2013). Interestingly, the authors observed no peaks and no differences in CKX activity at the particular stages of spike development, which is in agreement with homeostasis of the pool of isozymes in 7 DAP spikes of wheat, as suggested by us, which is independent of the level of silencing of *TaCKX1* but is rather a consequence of co-regulation of expression of other *TaCKX* genes. A similar effect of increased grain yield, which was a consequence of higher spike and grain number, was obtained in barley with silenced by RNAi *HvCKX1*, an orthologue of *TaCKX1* (Zalewski et al., 2010; Zalewski et al., 2014; Holubova et al., 2018). The three CKs measured, cZ, tZ and iP, were differently regulated in earlier or later, but not precisely defined, stages of spike development (Holubova et al., 2018); therefore the results of CK contents are not comparable. Incomparable to the results obtained for RNAi silenced *TaCKX1* and *HvCKX1*, no changes in yield parameters, including spike and seed number, were observed in mutant lines with knock-out of *HvCKX1* (Gasparis et al., 2019). These essential phenotypic differences between RNAi-silenced *TaCKX1* and *HvCKX1* or *HvCKX1* knocked out by CRISPR-Cas9 might be the result of different processes involved in inactivation of the gene. The first one is regulated at the posttranscriptional and the second at the transcriptional level. Since CKs might regulate various developmental and physiological processes at the posttranscriptional level (Cerny et al., 2011; Kim et al., 2012) or by modulation of context-dependent chromatin accessibility (Potter et al., 2018), the way of deactivating *TaCKX* function seemed to be important.

Spike number and grain number are highly correlated in both non-silent and silent plants and are regulated by the same groups of *TaCKX* genes as well as phytohormones. The first group includes *TaCKX1*, *2.2* and *5* positively regulating iP but negatively BA. The second comprises *TaCKX11 (3)* and *2.1* acting in the opposite way, and homeostasis of these hormones in non-silenced plants maintains a lower spike number. The main role in controlling higher spike and grain number in silent plants seemed to be played by *TaCKX2.1* and *5*. Their increased expression determine higher spike and grain number. The higher spike number was correlated with higher DZGs, iP and GA content. These correlations are not significant because they were measured in a stage of plant development in which the number of spikes and seed number have already been set.

As reported, the higher spike number was the consequence of a higher tiller number, which was positively correlated with the content of endogenous zeatin in the field-grown wheat after exogenous hormonal application (Cai et al., 2018). Moreover, the authors showed that IAA application inhibited the occurrence of tillers, by changing the ratios of IAA and ABA to zeatin. Shoot branching might also be dependent on acropetal transport of cytokinin (Shimizu-Sato et al., 2009).

Root weight was positively correlated with *TaCKX9 (10)* expression in 7 DAP spikes of non-silent plants and, conversely, negatively regulated by increased expression of this gene in silenced plants. Therefore lower (compared to silent plants) expression of the gene in the investigated organ determined lower root weight in the first group of plants, but higher in the second. *TaCKX9 (10)*, which is highly and specifically expressed in leaves (Ogonowska et al. 2019) and showed increased expression in 7 DAP spikes of silent plants, down-regulated cZOG. The same cZOG was up-regulated by *TaCKX11 (3)*, but expression of this gene in 7 DAP spikes of silent plants is strongly decreased. Both cZ and cZOG are involved in spike length regulation as well as TGW and grain yield in the group of silenced plants.

Since our results are restricted to the 7 DAP spike but are correlated with weight of seedling roots, we should take into consideration possible action of cytokinin transport and signalling genes as well as other phytohormones, which take part in hormonal cross-talk to control regulation of root growth (Pacifici et al., 2015). Accordingly cZ type CKs found as the major forms in phloem and are translocated from shoots to roots (Corbesier et al., 2003; Hirose et al., 2008). It is also worth underlining that only *TaCKX2.1* and *2.2* are specifically expressed in developing spikes. Both *TaCKX9 (10)*, highly and specifically expressed in leaves, and *TaCKX11 (3)*, expressed in all organs, seemed to regulate seedling roots as well, although in the opposite manner.

The lower plant height and higher root weight observed in the group of silenced plants of wheat is in agreement with opposed regulation of these traits by CKs and IAA mentioned above (Brenner and Schmulling 2012). CKs are considered as negative regulators of root growth, so their reduction causes enhanced root growth (Werner et al., 2010; Mrizova et al., 2013), while excess of CKs inhibits primary root growth in *Arabidopsis* (Riefler et al., 2006; Dello Ioio et al., 2012). It was also demonstrated that long-distance, basipetal transport of cytokinin controls polar auxin transport and maintains the vascular pattern in the root meristem (Bishopp et al., 2011). Therefore CKs might operate differently in distinct parts of the plant. The cZOG, represented mainly by inactive cZ-0-glucoside, down-regulated by *TaCKX9* (*10*) in 7 DAP spikes, was probably up-regulated in seedling roots, resulting in higher root weight. Content of cZ riboside was proved to be most abundant in the roots of maize (Veach et al., 2003). Opposite data, up-regulated content of active cZ in 7 DAP spikes, might influence down-regulation of this CK in roots. It has been documented that such suppressing cZ levels mediated by overexpression of *AtCKX7* affected root development in *Arabidopsis* (Kollmer et al., 2014). Higher weight of seedling root was also obtained by silencing via RNAi or knock-out via CRISPR/Cas9 of *HvCKX1* in barley plants, as in wheat, and the trait corresponded with decreased activity of CKX enzyme measured in roots (Zalewski et al. 2010, Gasparis et al. 2019).

Some *CKX* genes might be induced by transcription factors (Reid et al., 2016; Mao et al., 2019). An important role of some NAC transcription factors in regulation of *TaCKX* expression and subsequent organ development has also been documented in our not yet published results. Possible mechanisms of co-regulation of all these genes in the developing roots of silent and non-silent plants is being further investigated by us.

Leaf senescence was determined in the flag leaf of the first spike by measuring chlorophyll content. The trait was down-regulated by increased *TaCKX2.1* expression in silent plants. Opposite to the general role of *TaCKX* action, increased expression of the gene does not down-regulate but up-regulates tZ, tZ derivatives and cZ content in 7 DAP spikes. *TaCKX2.1* functions in a similar way, by up-regulating tZ, tZ derivatives and cZ in determining lower TGW and higher grain yield in silent T_2_ plants. Lower content of active CKs in the first spikes of non-silent plants is expected to up-regulate these CKs in flag leaves of these spikes, maintaining prolonged chlorophyll content. In contrast, higher content of tZ, tZGs, cZ and DZGs as well as GA in 7 DAP spikes of silent plants is expected to down-regulate CKs in the flag leaves, accelerating their senescence, which is documented by the results.

It was previously demonstrated that level of chlorophyll content in flag leaves is associated with the senescence process, in which CKs suppress inhibition of senescence (Gan and Amasino, 1995). During this processes, proteins are degraded and nutrients are re-mobilised from senescing leaves especially to the developing grains (Gregersen et al., 2008). We might suppose that slower spike ripening in non-silent plants, which is dependent on lower CK content in the 7 DAP spike, causes slower flow of micronutrients as well as CKs from flag leaf to spike. Therefore prolonged chlorophyll content in the flag leaf of the first spike negatively correlated with TGW but positively with plant height. Opposite data were obtained for flag leaves of silent plants, in which higher content of CKs in 7 DAP spikes might be the result of faster flow accelerating leaf senescence. Reduced chlorophyll content in flag leaves of the first spike of silent plants positively correlated with grain yield.

The important role of tZ and less active cZ in suppression of senescence was proved in maize leaves (Behr et al., 2012), and in an oat-leaf assay (Gajdosova et al., 2011). It was also documented that delayed senescence of wheat stay-green mutant, tasg1, at the late filling stage was related to high cytokinin and nitrogen contents (Wang et al., 2019).

## CONCLUSION

Based on the 7 DAP spike as a research object, we have documented that silencing of *TaCKX1* by RNAi strongly influenced up- or down-regulation of other *TaCKX* genes, as well as phytohormone levels and consequently phenotype. This co-regulation is dependent on the level of silencing of the gene and is independent of cross-silencing of other *TaCKX*.

In the general model of regulation of yield-related traits, highly silenced *TaCKX1*, which is specific for developing spikes, strongly down-regulates *TaCKX11 (3)*, expressed through different organs of the developing plant. Coordinated action of these genes is expected to lead to increased contents of tZ, tZGs, cZ and DZGs as well as GA, determining a high-yielding phenotype. In contrast, up-regulated *TaCKX2.1*, which is specifically expressed in developing spikes, and *TaCKX9 (10)*, strongly expressed in leaves, might be down-regulators of cZOG. However, detailed analysis revealed that each tested yield-related trait is regulated by various up- or down-regulated *TaCKX* genes and phytohormones. Key genes involved in regulation of grain yield, TGW or root weight in highly silenced plants are *TaCKX2.1* and *TaCKX11 (3)* acting antagonistically; increased expression of the first one determines growth of tZ, tZ derivatives and cZ, whereas decreased expression of the second down-regulates content of cZOG. A key role in determination of the high-yielding phenotype seemed to be played by the growing content of tZ in 7 DAP spikes, which might accelerate maturation of immature grains by speeding up nutrient flow from flag leaves. This finally led to reduction of TGW but enhancement of grain number and yield. The latter traits are the result of a higher spike number, which is determined in the early stages of plant development.

## Materials and methods

### Vector construction

The hpRNA type of silencing cassette was constructed in pBract207 (https://www.jic.ac.uk/technologies/crop-transformation-bract/). It contains the *Hpt* selection gene under the 35S promoter and cloning sites for the cloning silencing cassette under the Ubi promoter. The vector is compatible with the Gateway cloning system. For cloning purposes a coding sequence of *TaCKX1* (NCBI JN128583) 378 codons long was used. In the first step, the cassette was amplified using: EAC11-F: 5′-TTGAATTCGACTTCGACCGCGGCGTTTT-3′ and EAC12-R: 5′-TTGAATTCATGTCTTGGCCAGGGGAGAG-3 and cloned into the entry vector pCR8/GW/TOPO (Invitrogen). In the next step, the cassette was cloned to the destination Bract7 vector in the Gateway reaction. The presence of the silencing cassette in the vector was verified by restriction analysis and sequencing. The vector was electroporated into the AGL1 strain of *Agrobacterium tumefaciens* and used for transformation.

### Plant material, *Agrobacterium*-mediated transformation and *in-vitro* culture

The spring cultivar of common wheat (*Triticum aestivum* L.) Kontesa was used as a donor plant for transformation experiments as well as transgenic plants. Seeds were germinated into Petri dishes for one day at 4°C and then five days at room temperature in the dark. Six out of ten seedlings from each Petri dish were replanted into pots with soil. The plants were grown in a growth chamber under controlled environmental conditions with 20°C/18°C day/night temperatures and a 16 h light/8 h dark photoperiod. The light intensity was 350 µmol· s^-1^·m^-2^. *Agrobacterium*-mediated transformation experiments were performed according to our previously described protocols for wheat (Przetakiewicz et al., 2003; Przetakiewicz et al., 2004). Putative transgenic plants were regenerated and selected on modified MS media containing 25 mg l^-1^ of hygromycin as a selectable agent.

First, 7 days after pollination (DAP) spikes from T_1_, T_2_ and control plants were collected for RT-qPCR and phytohormone quantification. Only 1/3 of the middle part of each spike was used for experiments (upper and lower parts were removed).

### PCR analysis

Genomic DNA was isolated from well-developed leaves of 14-day plants according to the modified CTAB procedure (Murray and Thompson, 1980) or by using the KAPA3G Plant PCR Kit (Kapa Biosystems). The PCR for genomic DNA isolated by CTAB was carried out in a 25 ml reaction mixture using Platinum Taq DNA Polymerase (Invitrogen) and 120 ng of template DNA. The reaction was run using the following program: initial denaturation step at 94°C for 2 min, 35 cycles of amplification at 94°C for 30 s, 65°C for 30 s, 72°C for 30 s with a final extension step at 72°C for 5 min. The PCR for genomic DNA isolated by KAPA3G was carried out in a 50 μl reaction mixture using 1 U of KAPA3G Plant DNA Polymerase and a 0.5 x 0.5 mm leaf fragment. The reaction was run using the following program: initial denaturation step at 95°C for 3 min, 40 cycles of amplification at 95°C for 20 s, 68°C for 30 s, 72°C for 30s with a final extension step at 72°C for 2 min.

Putative transgenic T_0_ and T_1_ plants were tested with two pairs of specific primers amplifying a fragment of the *hpt* selection gene. The sequences of the primers for the first pair were: hygF1 5′-ATGACGCACAATCCCACTATCCT-3′ and hygR1 5′-AGTTCGGTTTCAGGCAGGTCTT-3′, and the amplified fragment was 405 bp. The sequences of the primers for the second pair were: hygF2 5′-GACGGCAATTTCGATGATG-3′ and hygR2 5′-CCGGTCGGCATCTACTCTAT-3′, and the amplified fragment was 205 bp. Non-transgenic, null segregants were used as a control.

### RNA extraction and cDNA synthesis

Total RNA from 7 DAP spikes was extracted using TRI Reagent and 1-bromo-3-chloropropane (BCP) (Sigma-Aldrich) according to the manufacturer’s protocol. The purity and concentration of the isolated RNA were determined using a NanoDrop spectrophotometer (NanoDrop ND-1000) and the integrity was checked by electrophoresis on 1.5% (w/v) agarose gels. To remove the residual DNA the RNA samples were treated with DNase I, RNase-free (Thermo Fisher Scientific). Each time 1 µg of good quality RNA was used for cDNA synthesis using the RevertAid First Strand cDNA Synthesis Kit (Thermo Fisher Scientific) following the manufacturer’s instructions. The obtained cDNA was diluted 20 times before use in RT-qPCR assays.

#### Quantitative RT-qPCR

RT-qPCR assays were performed for 6 target genes: *TaCKX1* (JN128583), *TaCKX2.1* (JF293079)/*2.2* (FJ648070), *TaCKX11* (*3*) (JN128585), *TaCKX5* (Lei et al. 2008), *TaCKX9* (*10*) (JN128591). Primer sequences designed for each gene as well as for the reference gene are shown in **Table S1**. All real-time reactions were performed in a Rotor-Gene Q (Qiagen) thermal cycler using 1x HOT FIREPol EvaGreen qPCR Mix Plus (Solis BioDyne), 0.2 µM of each primer, and 4 µl of 20 times diluted cDNA in a total volume of 10 µl. Each reaction was carried out in 3 technical replicates at the following temperature profile: 95°C – 15 min initial denaturation and polymerase activation (95°C – 25 s, 62°C – 25 s, 72°C – 25 s) x 45 cycles, 72°C – 5 min, with the melting curve at 72–99°C, 5 s per step. The expression of *TaCKX* genes was calculated according to the two standard curves method using *ADP-ribosylation factor* (*Ref 2*) as a normalizer.

Relative expression/silencing of *TaCKX1* was related to mean expression of the gene in non-silenced control plants set as 1.00. Relative expression of other *TaCKX* genes was related to each tested gene set as 1.00 in non-silenced plants.

Statistical analysis was performed using Statistica 13 (StatSoft) software. The normality of data distribution was tested using the Shapiro-Wilk test. To determine whether the means of two sets of data of expression levels, phytohormone concentrations, and yield-related traits between non-silenced and silenced lines are significantly different from each other (for p value less than p<0.05) Student’s t-test or the Mann-Whitney test was applied. Correlation coefficients were determined using parametric correlation matrices (Pearson’s test) or a nonparametric correlation (Spearman’s test).

### Quantification of ABA, auxins, cytokinins and GA_3_

Chemicals used for quantification were: the standard of ABA, five standards of auxins: IAA, indole-3-butyric acid (IBA), indole-3-propionic acid (IPA), 1-naphthaleneacetic acid (NAA), and 2-phenylacetic acid (PAA); twenty-seven standards of CKs: *t*Z, *trans*-zeatin riboside (*t*ZR), *trans*-zeatin-9-glucoside (*t*Z9G), *trans*-zeatin-7-glucoside (*t*Z7G), *trans*-zeatin-*O*-glucoside (*t*ZOG), *trans*-zeatin riboside-*O*-glucoside (*t*ZROG), *trans*-zeatin-*9*-glucoside-*O*-glucoside (*t*Z9GOG), *trans*-zeatin-9-glucoside riboside (tZ9GR), *c*Z, *cis*-zeatin-riboside (*c*ZR), *cis*-zeatin *O*-glucoside (*c*ZOG), *cis*-zeatin 9-glucoside (*c*Z9G), *cis*-zeatin-*O*-glucoside-riboside (*c*ZROG), dihydrozeatin (DZ), dihydrozeatin-riboside (DZR), dihydrozeatin-9-glucoside (DZ9G), dihydrozeatin-7-glucoside (DZ7G), dihydrozeatin-*O*-glucoside (DZOG), dihydrozeatin riboside-*O*-glucoside (DZROG), *N*^6^-isopentenyladenine (iP), *N*^6^-isopentenyladenosine (iPR), *N*^6^-isopentenyladenosine-7-glucoside (iP7G), *para*-topolin (*p*T), *meta*-topolin (*m*T), *ortho*-topolin (*o*T), 6-benzylaminopurine (6-BAP), and standard of GA_3_ were purchased from OlChemIm (Olomouc, Czech Republic). Methanol (MeOH), acetonitrile (ACN), water (LC-MS purity), and formic acid (FA) were purchased from Merck KGaA (Darmstadt, Germany).

For the measurement of phytohormones, 200 mg of plant powders were placed into the 2 mL Eppendorf tubes, suspended in 1 mL of (v/v) 50% ACN and homogenized in a bead mill (50 Hz, 5 min; TissueLyser LT, Qiagen, Germany) using two 5 mm tungsten balls. Then, samples were homogenized using the ultrasound processor VCX 130 (max. power 130 W, max. frequency 20 kHz, 5 min) equipped with titanium probe (Sonics & Materials Inc., USA) and mixed in laboratory shaker (90 rpm, dark, 5°C, 30 min; LC-350, Pol-Eko-Aparatura, Poland). Samples were centrifuged (9000×g, 5 min; MPW-55 Med. Instruments, Poland) and collected in a glass tube. For quantification of ABA, AXs, CKs and GA_3_, [^2^H_6_](+)-*cis*,*trans*-ABA (50 ng), [^2^H_5_] IAA (15 ng), [^2^H_6_] iP (50 ng), [^2^H_5_] *t*Z (30 ng), [^2^H_5_]-*t*ZOG (30 ng), [^2^H_3_]-DZR (30 ng) and [^2^H_2_] GA_3_ (30 ng) were added to samples as internal standards.

Prepared extracts were purged using a Waters SPE Oasis HLB cartridge, previously activated and equilibrated using 1 mL of 100% MeOH, 1 mL water, and 1 mL of (v/v) 50% ACN (Simura et al., 2018). Then, extracts were loaded and collected to the Eppendorf tubes and eluted with 1 mL of 30% ACN (v/v). Samples were evaporated to dryness by centrifugal vacuum concentrator (Eppendorf Concentrator Plus, Germany), dissolved in 50 µL of (v/v) 30% ACN and transferred into the insert vials. Detection of analyzed phytohormones was performed using an Agilent 1260 Infinity series HPLC system (Agilent Technologies, USA) including a Q-ToF LC/MS mass spectrometer with Dual AJS ESI source; 10 μL of each sample was injected on the Waters XSelect C_18_ column (250 mm × 3.0 mm, 5 μm), heated up to 50 °C. Mobile phase A was 0.01% (v/v) FA in ACN and phase B 0.01% (v/v) FA in water; flow was 0.5 mL min^-1^. Separation of above hormones was done in ESI-positive mode with the following gradient: 0-8 min flowing increased linearly from 5 to 30% A, 8-25 min 80% A, 25-28 min 100% A, 28-30 min 5% A.

For the optimization of MS/MS conditions, the chemical standards of analysed phytohormones were directly injected to the MS in positive ([M + H]^+^) ion scan modes, then areas of detected standard peaks were calculated. [M + H]^+^ was chosen because of its significantly better signal-to-noise ratios compared to the negative ion scan modes.

Chlorophyll content was measured using a SPAD chlorophyll meter.

## SUPPLEMENTAL DATA

Supplemental Table S1 Primer sequences designed for reference gene and each of 6 tested *TaCKX* genes and amplicon length.

Supplemental Table S2 Phenotypic traits and ratio indicator in silent T_1_ and not silent, control plants.

Supplemental Table S3 Phenotypic traits and ratio indicator in silent T_2_ and not silent, control plants.

Supplemental Table S4 A. B. Correlation coefficients among expression of all tested *TaCKX* genes and enzyme activity, and phenotypic traits in non-silent (A) and highly silent T_2_ plants (B). * non-parametric analysis; in bold - significant at p<0.01.

## ACKNOWLEDGEMENTS

This research was supported by the National Science Centre, grant UMO-2014/13/B/NZ9/02376 and a statutory grant of PBAI-NRI.

We thank Malgorzata Wojciechowska, Izabela Skuza, Agnieszka Glowacka, Dr. Maja Boczkowska and Agnieszka Onysk for excellent technical assistance.

